# Repetitive execution of a reach-and-lift task is associated with inter-trial changes in movement-related EEG features and their predictive power

**DOI:** 10.1101/2023.02.09.527923

**Authors:** Andrew Paek, Shikha Prashad

## Abstract

Brain Machine Interfaces (BMIs) can help restore motor function to individuals with paralysis. These systems allow users to control an assistive device through the detection of movement-related brain activity. Such neural signatures are found through machine learning algorithms and training datasets that are generated from participants performing repetitive motor tasks. We anticipate that the movement-related brain waves of interest can attenuate over time due to neural efficiency, where the brain becomes more efficient with practice in a motor task. To explore this hypothesis, we used three open-access EEG datasets where participants performed a simple reach-and-lift task. From each trial, time windows associated with resting and movement periods were segmented. Alpha- and beta-band spectral power was estimated for each epoch, and event-related desynchronization (ERD) was estimated as the suppression in spectral power from rest to movement. These ERDs were compared between early and late trials in the dataset. We also used linear discriminant analysis to assess a machine learning algorithm’s accuracy in classifying whether the time windows belonged to rest or movement based on spectral power. In some cases, the ERDs were significantly different between earlier and later trials, and these differences led to changes in predicting the presence of movement from these ERDs. These results call for a reevaluation of BMI performance in datasets with numerous trials and an exploration of strategies that can compensate for longitudinal changes in movement-related brain activity used for BMIs.

## Introduction

Brain-machine interfaces (BMIs) are systems that allow users to control external devices through their thoughts^1–3^. These external devices can include assistive devices such as prostheses, powered exoskeletons, or robotic arms, all of which can be used to help restore motor function to an individual with paralysis. The key components in BMIs are the neuroimaging modality used to monitor a user’s brain activity and the machine learning algorithms that are used to detect neural patterns of interest to control an external device. These algorithms find a mapping between the variations in brain activity with different commands that would control the assistive device.

While a variety of neuroimaging techniques can be used for BMIs, scalp electroencephalography (EEG) remains the most popular due to its accessibility and noninvasive nature. A popular brain wave pattern that is used in EEG-based BMIs is event-related desynchronization (ERD)^4–8^. The ERD manifests as an attenuation in alpha- (8-13 Hz) or beta- (20-30 Hz) band power when the user is engaged in limb movement. The suppression is typically found in sensors that are close to the somatotopic cortical areas associated with the engaged limb^5,7–10^. For example, when the participant moves their right hand, a suppression in alpha-band power is found in the left sensorimotor areas associated with the hand regions. Based on the limb that is engaged and what areas of the brain are monitored, this pattern can be used to infer when the participant is moving and what limb is being engaged. This signature is also manifested during motor imagery, where the participant imagines moving their limb without overtly performing the limb movements^7,10,11^. This phenomenon makes the pattern favorable for BMIs since individuals with paralysis can perform the motor imagery to modulate these movement-related ERDs^12,13^.

From this movement-related ERD, a BMI detects when a user is engaged with limb movements through a programmed predictive model. It is possible to create a predictive model that monitors the spectral power from a few sensorimotor sensors (e.g., C3 and C4), and manually tunes thresholds that correspond to motor commands of interest. However, it is conventional to generate predictive models through machine learning algorithms^14–16^. These algorithms allow predictive models to incorporate many features, which typically increases a predictive model’s accuracy. Many EEG-based BMIs incorporate features from multiple EEG sensors and multiple features (e.g., power from alpha- and beta-bands) from each EEG sensor. In addition, machine learning algorithms can better tune a predictive model’s parameters to better match each individual user. This need for personalization can be due to idiosyncratic variations in the participant’s brain activity^16^, where it is unknown which sensors will provide the most optimal signal, and what thresholds should be used with the spectral power estimates to distinguish when the limb is engaged or idle.

To accommodate an individual’s unique brain waves, the machine learning algorithms are trained based on the individual’s dataset that contains examples of the brain waves with the associated motor commands of interest. This dataset is typically created as the participant performs a motor task while the participant’s neural activity is recorded. It is common for these datasets to contain a large number of trials that can range from 100 – 600 trials ^17–23^, where the participant performs a motor task repetitively. A large number of trials helps highlight the movement-related cortical patterns of interest through grand averaging and allows the predictive models to be trained and validated through more data.

The conventional notion in BMI studies is that training a predictive model with more data will improve decoding performance. However, these algorithms rely on the assumption that the relationship between the predictive features and the command of interest is consistent throughout the trials. If the mapping changes throughout the dataset, it can cause the machine learning model to perform poorly^24^. We argue that such non-stationarities may be present in BMI datasets due to neural efficiency, a phenomenon where the brain becomes more efficient in executing well-practiced skills. This phenomenon has been observed in neuroimaging studies where experts displayed less cortical activation compared to novices when both groups performed the same task^25–27^. Additionally, studies using transcranial magnetic stimulation have found that the motor cortex can undergo plasticity in 30-minute sessions of repeated movements^28,29^. We hypothesize that when participants in BMI studies perform a repetitive motor task, their movement-related brain waves of interest are likely to attenuate across trials within a dataset. Since these non-stationarities are more pronounced in studies with more trials, it is possible that BMI studies with numerous trials could yield lower accuracies than those with fewer trials. This association could complicate comparisons between different decoding strategies if the BMI studies used to benchmark them differ considerably in the number of trials. While previous studies have argued that ERDs are consistent across trials and sessions, the measures that are reported only show a moderate amount of consistency and can vary significantly depending on the behavioral task, sensor location, and parameters used to extract spectral power features^30–32^.

In this work, we explored whether numerous trials cause movement-related ERDs to attenuate over time in three open access datasets where EEG was collected while participants performed repetitive reach-and-lift tasks. We also examined whether this attenuation influences the predictive power of machine learning algorithms in classifying when participants are resting vs. moving. We hypothesize that movement-related ERDs will attenuate in later trials compared to earlier trials. We also hypothesize that classification accuracies will be higher when the dataset only includes the earlier trials, compared to when the dataset includes only the later trials and both early and later trials due to the changes in ERD strength.

## Methods

### Open access datasets

We used three open access EEG datasets in our study. These datasets were chosen based on the recording modality, behavioral task, and the number of trials. All datasets used EEG recordings performed with gel-based EEG caps. The behavioral task used real movements associated with reaching and grasping an object. All datasets had approximately 150 trials for each participant.

The first dataset was from Luciw and colleagues^17^ where 12 participants (all right-handed, 8 female participants, ages between 19-35 years, no commentary on the lack of neurological conditions) performed a behavioral task where they reached, grasped, and lifted a customized device. The device contained force and kinematic sensors, and a light-emitting diode (LED) that served as a visual cue. Participants were instructed to rest their hand in the starting position and start reaching for the grasped device when the LED turned on. After grasping the device, they lifted the device off the table until the LED turned off. This cued the participant to set down the device and return their hand to the starting position. Each participant performed 10 blocks, each of which contained 28-32 trials. Each block contained experimental variations where the experimenter changed the device’s weight and/or the grip surface. The experimenter’s changes in the device’s properties were hidden from the participant. A 32-channel 500 Hz EEG system (ActiCap electrodes with BrainAmp EEG signal amplifier, Brain Products GmbH, Gilching, Germany) was used to record brain activity, five electromyography (EMG) sensors were used to record muscular activity around the hand and arm, and kinematic and kinetic sensors associated with the hand and arm were recorded.

The second dataset was from Schwarz et al.^33,34^, where 15 participants (all right handed, 5 female participants, ages between 15 and 30, no known medical conditions) performed a behavioral task that involved reaching and grasping one of two objects that are commonly used in daily living. The objects were designed such that one required a palmar grasp while the other required a lateral grasp. Participants were instructed to rest their palm on the starting location, which contained a force sensor that tracked the onset of movement. Each trial was self-initiated, where participants were instructed to focus on one of two objects for 1-2 seconds and then reach and grasp one of the objects for 1-2 seconds. Participants rested for four seconds in between trials. From each participant, we extracted 139 to 160 trials, based on the event markers for the onset of movement. For their study associated with the open-access dataset, comparisons were made between a gel-based, water-based, and dry EEG system recordings^33^. We also note that the dataset was part of another study that included bimanual movements, but the shared dataset only includes trials with unimanual movements^34^. For our study, we only used the data collected with the gel-based hardware. A 64-channel 256 Hz system (g.USBamp and g.GAMMAsys/g.LADYbird active electrode system, g.tec medical engineering GmBH, Austria) was used to record brain activity, six of which were used to record electrooculograms (EOG). Force resistive sensors were used to monitor the movement onset and when the object was grasped.

The last dataset was from Jeong and colleagues^20^ where 25 participants (all right-handed, 10 female participants, ages between 24-32 years, no known neurophysiological anomalies or musculoskeletal disorders) performed a variety of upper limb movement tasks that included reaching in eight different directions, reaching and grasping three different objects, and twisting the wrist clockwise and counterclockwise. Participants performed all three tasks with real and imaginary movements in three single-day sessions that were at least one week apart. For our study, we only included data from the first session that corresponded to the grasping task. For this task, participants were instructed to reach and grasp one of three objects that were common in daily living. Each object required a different grasp shape that was cylindrical, spherical, or lateral. Participants performed the task based on visual cues shown on a computer monitor: a 3-second rest cue where participants relaxed, followed by a 4-second preparatory cue that instructed the participant which object to grasp next, then followed by a 4-second movement cue that indicated when participants should start moving. The dataset contained exactly 150 trials for each participant. A 64-channel 2500 Hz EEG system (BrainAmp, Brain Products GmBh, Germany) was used to record brain activity, four of which were used to record EOG. To help reduce the computational time for our study, these signals were resampled to a rate of 1000 Hz.

### EEG preprocessing

EEG signals were preprocessed to help reduce the artifacts in the data. To remove the drift in the signals, we used a high-pass filter with a zero-phase 4^th^ order Butterworth filter at a cutoff of 0.1 Hz. To remove brief deflections in the signals likely caused by head or sensor movements, artifact subspace reconstruction (ASR) was performed^35,36^. Time windows with a length of 0.5 seconds that had a variance beyond 60 standard deviations compared to quiescent parts of the data were corrected with ASR. Lastly, independent component analysis (ICA) was used to find components associated with ocular, muscular, cardiac, and line noise activity^37^. These were automatically detected and removed with the “IClabel” functions, which classifies components based on a pre-made database^38^. ASR, ICA, and IClabel were performed with the EEGLAB toolbox^39^ in MATLAB (Mathworks, Natick, Massachusetts, USA) ^40^. While EOG sensors were used in ICA to reduce artifacts, they were omitted for the rest of the analysis.

### Comparing ERDs between early and late trials

From each trial, a 1-second time window was extracted during the rest and movement periods. These periods were extracted based on the event markers that were present in the datasets. For Luciw et al.’s and Schwarz et al.’s datasets, we utilized the event markers associated with the onset of reach provided by the authors. For Jeong et. al.’s dataset, we used the onset of the visual cue that instructed when participants would start the movement. The rest period was extracted from the trial -2 to -1 seconds with respect to the subsequent movement onset. The movement period was extracted from the trial as the 1-second epoch immediately after movement onset.

For each of these rest and movement periods, spectral power was extracted based on the power spectral density calculated using the Welch method. Within the one-second time period, the averaged Fast Fourier Transform solution used sample windows with a length of 0.5 seconds and an overlap of 0.25 seconds. The Hamming window was used to reduce spectral leakage. From the power spectral density (PSD), spectral band power was extracted in the alpha (8-13 Hz) and beta (20-30 Hz) bands.

For each trial, a single value of the alpha- and beta-band power was extracted for each rest and movement period. For each participant, the spectral power values were aggregated from the trials that belonged to the early and late portions of the session, each of which contained approximately 50 trials. Since these are time periods of interest in exploring the effects of task repetition, we designate these portions as early and late “trial-groups” for the rest of the paper. For Luciw et al.’s dataset, the session was divided into 10 blocks of trials which contained different experimental changes to the object’s properties. We used blocks 1, 2, and 3 as the early trial-group and blocks 7, 8, and 9 as the late trial-group. Since the authors intentionally omitted kinematic information in the 10^th^ block of trials, it was omitted in our study. For Schwarz et al.’s ^33^ dataset, each participant had a different number of trials, so the number of trials was divided into thirds, and the first and last third groups of trials were respectively designated as the early and late trial-groups. For Jeong et al.’s^20^ dataset, each participant performed exactly 150 trials so the first and last 50 trials were respectively designated as the early and late trial-groups. A generic example of these early and late trial-groups is shown in Figure 1A.

**Figure 1).**
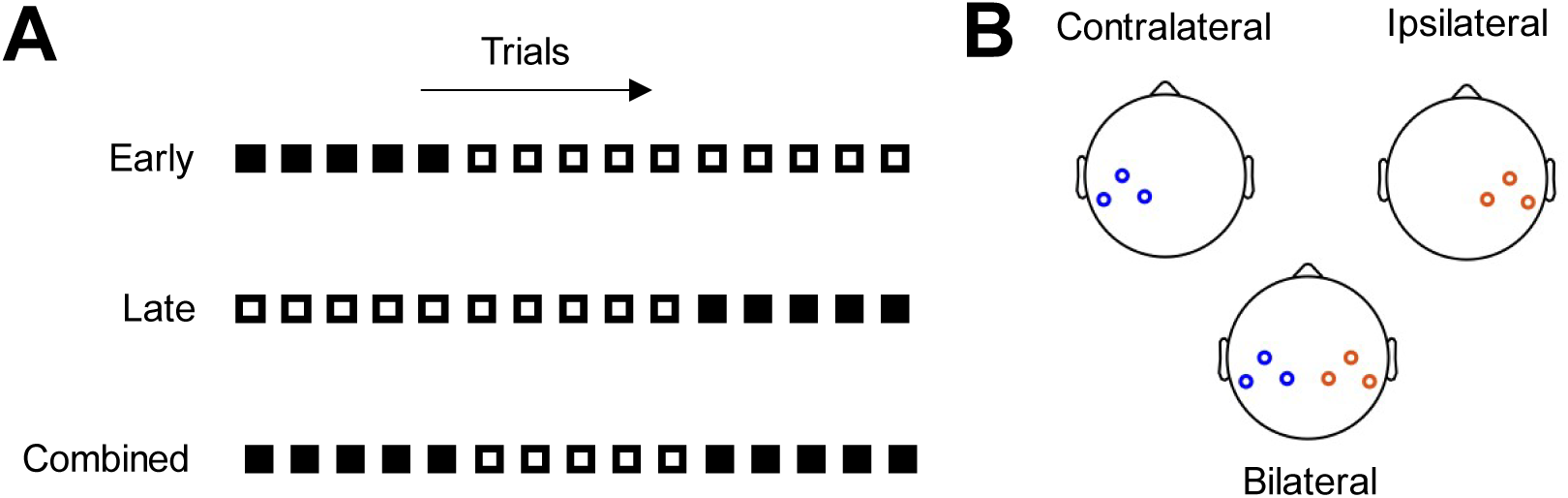
Diagrams indicating trial-groups and channel-groups used for comparisons. A) The groups of trials in a session that were designated as early, late, and combined trial-groups. The squares shown in black illustrate which trials were included for each trial-group. Each trial-group consisted of approximately 50 trials from each participant in each dataset. B) These scalp maps indicate the group of channels used in the decoding analysis, i.e., sensors that were contralateral to the hand (C3, CP5, CP1), ipsilateral to the hand (C4, CP2, CP6), or bilateral (all six sensors).

Alpha- and beta-band ERDs were compared between early and late trial-groups. For each trial, the ERD was calculated as the base-10 logarithm of the ratio in the spectral power during the movement period over the spectral power during the resting period. For each participant, the median of the ERDs was calculated across all the trials. A paired sample t-test was performed to compare the ERD values between the early and late trial-groups across all participants. This process was performed on each EEG channel. To help visualize the spatial distribution of the ERD values, they were also plotted as a scalp map.

### Decoding rest vs. movements from EEG

We also wanted to explore if the possible attenuation of the ERDs influences the machine learning model’s ability to predict when participants were moving or resting their hands. To test this, we designed a classification problem where we classified if participants were moving or resting during the one-second time periods that were segmented from each trial. The predictive features were the base-10 logarithm of the alpha- and beta-band power values that were extracted as described in the previous section.

Linear discriminant analysis (LDA) was used to classify if the time window belonged to a period of rest or movement. This was trained in a 5-fold cross-validation. The cross-validation was performed under two permutations: by channel-groups and by trial-groups. For the channel groups, cross-validations were performed with channels contralateral to the hand (sensors C3, CP5, CP1), ipsilateral to the hand (C4, CP2, CP6), and with both groups of channels. These sensors were chosen because they were present in all three datasets and scalp maps of the ERDs showed localized strength in the central-parietal regions. For the trial-groups, cross-validations were performed with trials only in the early trial-group, the late trial-group, and both the early and late trial-groups combined. Figure 1 illustrates the permutations of the cross-validations used. For each participant and the permutation, the average of the classification accuracies was calculated across the folds. A paired sample t-test was used to compare classification accuracies for the participants between the permutations that involved early vs. late trial-groups. We also compared classifications for each condition between alpha and beta-band power and between channel groups.

## Results

### Spectral power modulations associated with movement

In the comparisons of ERDs between the early and late trial-groups, we found that the longitudinal differences varied between the three datasets. The scalp maps depicting the ERDs in the early and late trial-groups are shown in Figure 2.

**Figure 2.**
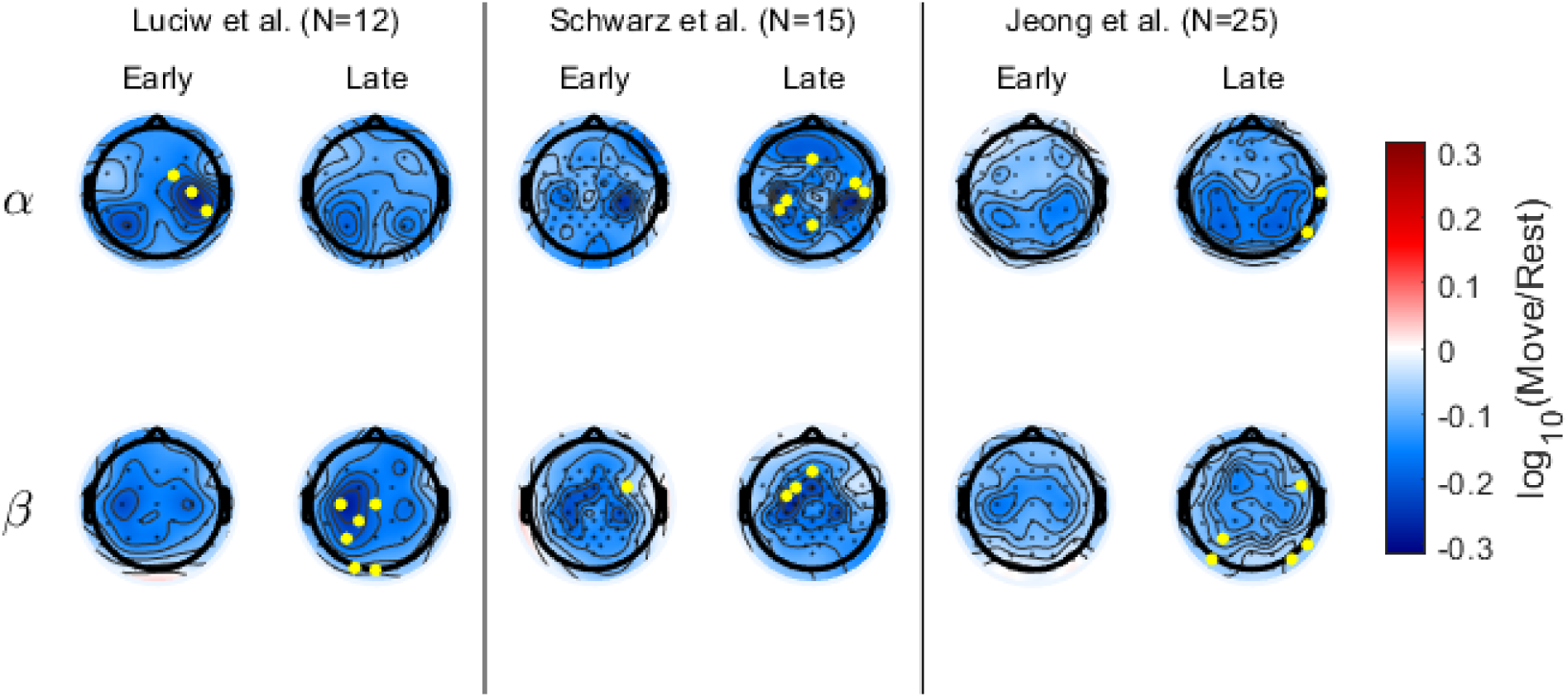
Scalp map of spectral power ERDs due to movement. The top and bottom rows correspond to alpha- (8-13 Hz) and beta-band (20-30 Hz) ERDs, respectively. For each dataset, the left and right columns correspond to ERDs from early and late trial-groups. Darker blue colors indicate stronger ERDs. A channel with a yellow circle on the early column indicates when the early trial-group had stronger ERDs compared to the late trial-group, and vice versa. (p<0.05).

In Luciw et al.’s dataset, the early trial-group had significantly stronger alpha-band ERDs in the ipsilateral central-parietal areas (FC2: p=0.020, C4: p=0.012, CP6, p=0.0066) while the late trial-group had significantly stronger beta-band ERDs in the contralateral central-parietal areas (C3: p=0.023, Cz: p=0.025, CP1: p=0.0064, P3: p=0.023, O1: p=0.019, Oz: p=0.038). In Schwarz et al.’s dataset, broader areas of the scalp had significantly stronger ERDs in the late trial-group for the alpha-band (Fz, p=0.0080, FCC6h: p=0.014, C6: p=0.037, CCP3h: p=0.014, CP3: p=0.017, Pz: p=0.040). In the beta-band, the contralateral frontal-central area had significantly stronger ERDs in the late trial-group (Fz: p=0.017, FC1: p=0.0048, FCC3h: p=0.00070). One sensor had significantly stronger beta-band ERDs in the early trial-group (FC4: p=0.024). In Joeng et al.’s dataset, the later trial-group had few peripheral sensors with significantly stronger ERDs in the alpha- (T8: p=0.044, P8: p=0.024) and in the beta- (P3: p=0.023, PO7: p=0.031, FC6: p=0.043, P8: p=0.013, PO8: p=0.0089) bands.

### Neural decoding performance in classifying rest vs. movement

Figure 3 contains boxplots of the classification accuracies for predicting whether the time periods from each trial belonged to movement or rest from the spectral power features. Tables of the statistics in the classification accuracies as well as the p-values for all comparisons can be found in the Appendix.

**Figure 3).**
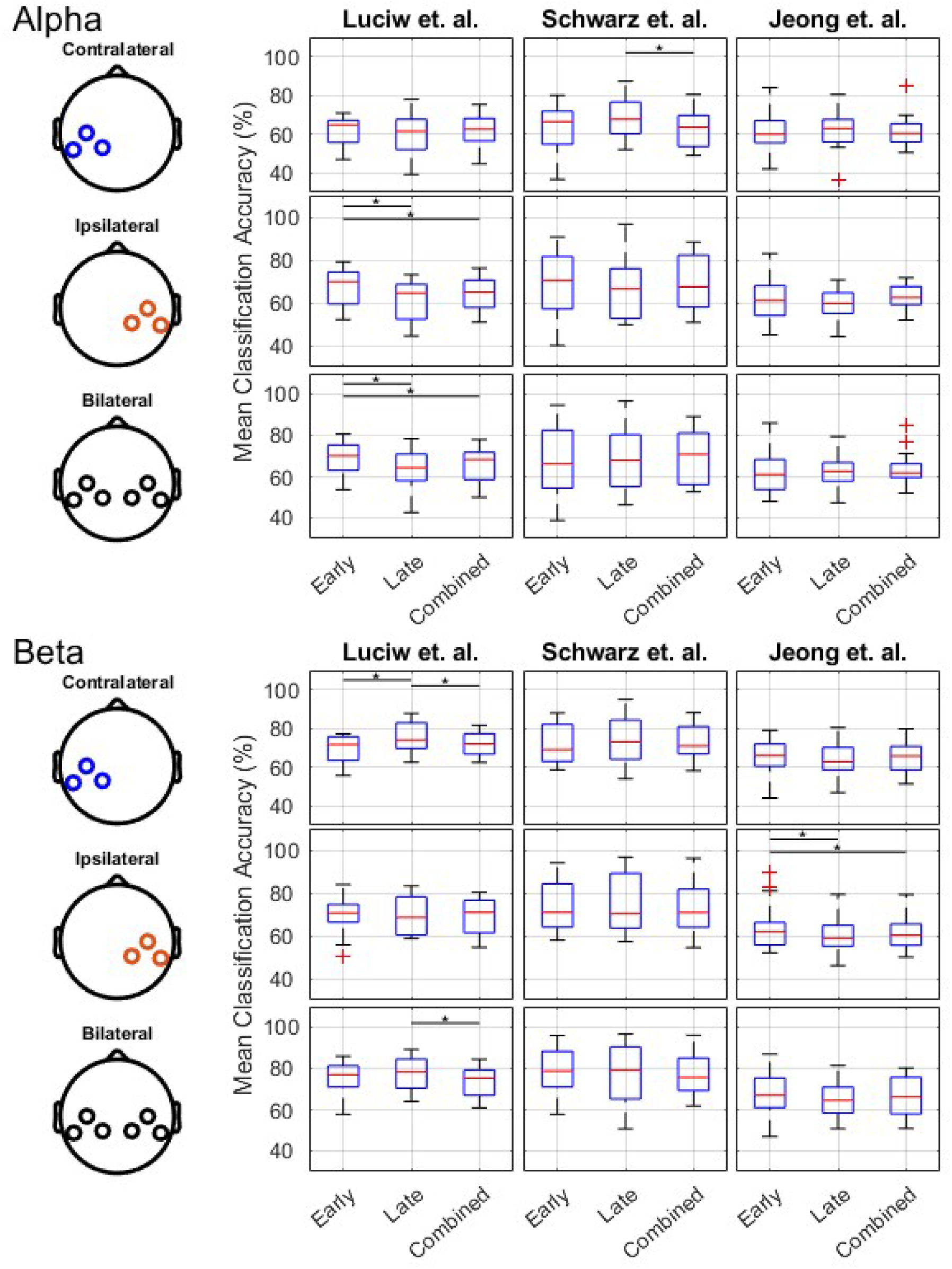
Boxplots of the classification accuracies for predicting rest vs movement with spectral band features. The top and bottom panels correspond to accuracies when alpha and beta-band power were used, respectively. Within each panel, each row corresponds to which group of channels were used in classification, and each column corresponds to what dataset was used. Each group of boxplots contains classifications that used trials from only the early trial-group (Early), only the late trial-group (Late), and from both trial-groups (Combined). Each boxplot corresponds to the spread of classification accuracies averaged across all cross-validation folds for each participant (N=12 for Luciw et al., N=15 for Schwarz et al., and N=25 for Jeong et al.). Asterisks indicate significant differences in classification accuracies between each condition based on paired sample t-tests (p<0.05).

Classifying with alpha-band power yielded significant differences between early and late trial-groups in Luciw et al.’s dataset and Schwarz et al.’s dataset. In Luciw et al.’s dataset with ipsilateral channels, the early trial-group yielded significantly larger accuracies than the late trial-group (p=0.0084) and combined early and late trial-groups (p=0.022). With the bilateral group of channels, the early trial-group yielded significantly larger accuracies than the late trial-group (p=0.032) and the combined early and late trial-group (p=0.042). In Schwarz et al.’s data, using the contralateral group of channels yielded significantly larger accuracies in the late trial-group compared to the combined early and late trial-groups (p=0.014).

Classifying with beta-band power yielded differences between early and late trial-groups in Luciw et al.’s and Jeong et al.’s datasets. In Luciw et al.’s dataset with contralateral channels, the late trial-group yielded significantly higher accuracies than the early trial-group (p=0.035) and the combined early and late trial-groups (p=0.012). Using both groups of channels also yielded significantly higher accuracies in the late trial-group compared to the combined early and late trial-group (p=0.033). In Jeong et al.’s dataset with ipsilateral channels, the early trial-group yielded significantly higher accuracies than the late trial-group (p=0.035) and the combined early and late trial-groups (p=0.018).

For most channel and trial-group conditions in Luciw et al.’s and Schwarz et al.’s datasets, beta-band power yielded significantly higher accuracies than alpha-band power (p <0.05; see appendix for all p-values). In comparisons in classification accuracies between channel groups (such as classifying with contralateral channels vs. ipsilateral channels), significant differences were found between channel groups, but they appeared to depend on what trials were used and changed depending on whether alpha- or beta-band power was used (see appendix for p-values).

## Discussion

In this work, we examined whether movement-related EEG features can attenuate over repetitive executions of a motor task. We also assessed if these changes affect a predictive model’s accuracy over time. Based on three open-access datasets where EEG was recorded during a reach-and-grasp task, we found different trends in how the spectral ERDs changed across trials, and their associated changes in classification performance.

### Changes in movement-related ERDs and their predictive power over time

In Luciw et al.’s dataset, we found that alpha-band ERDs became weaker over time in the ipsilateral areas while beta-band ERDs became stronger over time in the contralateral areas. This pattern coincided with the change in classification performance between early and late trials, where alpha-band power performed better in early trials while beta-band activity performed better in later trials. In Schwarz et al.’s dataset, we found that both alpha- and beta-band power had become stronger over time in a very sparse distribution across the scalp. This agrees with one classification case, where the contralateral group of channels in the late trials performed better than using the combined early and late trials. In Jeong et al.’s dataset, we found that a few sensors in the later trials had significantly stronger ERDs. However, this conversely coincided with a classification case where beta-band power from ipsilateral channels performed significantly better with early trials than using only later trials or combining both early and later trials.

We anticipated that task repetition would weaken movement-related ERDs over time due to the participant’s habituation to the task. This concept of neural efficiency has been demonstrated in experts, who displayed weaker neural modulations compared to novices when both groups performed the same task^25–27,41^. In addition, when participants were exposed to a repetitive stimulus, the event-related potentials associated with the stimulus became weaker over time^42,43^. Alpha-band modulations are also associated with cognitive load^44,45^, which we anticipated would reduce over time as participants became familiar with the task. EEG power and coherence with muscular electromyography activity have been shown to change with muscular fatigue^46–48^. We anticipated that such factors would weaken the movement-related ERDs over time as participants performed the same reach-and-grasp task over multiple trials. While we found significant changes between early and late trials in our study, we found some cases in classification and in the scalp maps where ERDs weakened and others where ERDs strengthened over time, depending on the channel group and the dataset.

We also explored classification accuracies between left and right central-parietal sensor groups since we anticipated that one side would be affected by task repetition. While it is generally understood that the brain hemispheres control contralateral sides of the limbs, it is not uncommon to observe bilateral activation during unimanual hand tasks. This is often seen as a bilateral alpha- and beta-band ERD during movements, with the contralateral ERD being stronger^7,8^. Given this laterality, we anticipated that the contralateral ERDs would be more affected by any inter-trial attenuation that would occur. Contrary to this hypothesis, however, we found that the ipsilateral ERDs can weaken over time, as shown in Luciw et al.’s alpha-band ERDs. We also found that later trials in Schwarz et al.’s and Jeong et al.’s datasets had stronger alpha- and beta-band ERDs in a sparse group of sensors. Additionally, we found that beta-band ERDs became stronger across trials in the contralateral side in Luciw et al.’s dataset. While we did not expect the beta-band ERD to strengthen, it further confirms that changes in ERDs across trials can be isolated in different spatial areas. This warrants consideration for BMIs that utilize features that incorporate the entire scalp (such as principal component analysis^49,50^ or common spatial patterns^51,52^). These techniques often apply weights across all sensors on the scalp and are fixed throughout the entire training dataset and may not accurately reflect spatially localized changes in ERDs.

Among the three datasets, Luciw et al.’s dataset demonstrated clearer changes in alpha- and beta-band ERDs over time and led to corresponding changes in classification performance. We suspect that this difference might be related to the varying novelty of the grasped object between the different datasets. In Schwarz et. al.’s^33^ and Jeong et al.’s^20^ datasets, participants were instructed to grasp objects that were typical in daily living. In Luciw et. al.’s^17^ study, participants grasped a novel cube-like device and the experimenter changed the grasped device’s weight and surface texture between trials without the participant’s knowledge. We suspect these experimental adjustments induced more cortical engagement associated with building internal models associated with the grasped device. Previous neuroimaging studies have highlighted the role of parietal regions in encoding physical and functional properties with grasped objects^53–56^. Motor learning studies have also found stronger beta-band suppression during periods of adaptation or sequence learning^57–60^. This may explain the stronger changes in ERDs we observed in Luciw et al.’s dataset, and not in the other datasets where object properties were fixed and more familiar to the participants.

### Implications for future brain machine interface and neuroimaging studies

In this work, we chose a simple scheme where we predicted the absence or presence of hand movements based on a reach-to-grasp task. It is uncertain whether differences in implementations of behavioral tasks and strategies used to predict cognitive states affect our findings and how these differences affect movement ERDs change over time. Many EEG-based BMIs are designed where multiple limbs are engaged by the participant, and the classifier is designed to distinguish which limb is moving^12,61–64^. While we chose alpha- and beta-band power as predictive features, other features such as readiness potentials or delta-band amplitude have also been used to predict movements, especially in reach-and-grasp tasks^65,66^. Lastly, many EEG-based BMI studies also use behavioral tasks that call for repetitive movements as opposed to short and brief movements like reaching and grasping^7,8,11,67,68^. Nonetheless, we chose a simple decoding scheme that utilizes alpha- and beta-band ERDs because this is a predictive feature that is very commonly used in many EEG-based BMIs. We also argue that these findings related to predicting rest vs. movement apply to other studies that predict more specific motor behaviors such as hand shapes^34,65,69,70^ or reach direction^70,71^. The findings in this work could potentially extend to other BMI control schemes and warrant consideration of how BMIs are evaluated in terms of accuracy.

We demonstrate that movement-related ERDs can significantly change over time, but we are unable to distinguish if the observed changes in ERDs could be attributed to factors associated with task habituation or fatigue. Distinguishing between these two factors could provide better guidance in deciding which strategies would yield more consistent neural modulations associated with movements. A strategy that can compensate for habituation would be omitting earlier trials from the analysis, since these are the time periods when participants are becoming familiar with the task. Another strategy that can compensate for fatigue would be including multiple sessions that have fewer trials or more breaks between blocks of trials. Tracking behavioral task performance during the trials could help distinguish these two types of factors, but this was absent in the datasets used in this study. BMI studies typically do not have a behavioral metric of performance since most incorporate a task of daily living that are well-practiced among participants. Future work that correlates task performance with the strength of ERDs across trials in a repetitive task could help distinguish these two factors, where a slow increase in task performance is likely to be associated with gradual skill acquisition.

In most cases where machine learning is used, predictive models usually become more accurate when they are trained with more data. In the case of BMIs, this convention generally leads researchers to collect more trials in a behavioral task with the expectation that a large dataset will yield high accuracies in the predictive models. However, the results of this study indicate that using a smaller subset of trials, particularly the earlier trials in session, could yield significantly larger accuracies than using all the trials that are available. This is likely to occur due to the non-stationarity of the predictive features throughout the trials in a session. When substantial non-stationarities are present, a predictive model is likely to have higher accuracies when they are cross-validated with fewer trials that cover a short time span. Longer non-stationarities would cause the model to perform worse on testing sets that cover a longer range in time. This “covariant shift”^72,73^ in the EEG features is well known and has motivated the development of adaptive algorithms that can retrain the predictive model over time^74–77^. To assess BMI feasibility studies and how decoder performance is reported, it warrants exploring if the predictive models might perform better with fewer trials than with the entire dataset. It also warrants considering the impact of the number of trials used in a given BMI feasibility study since the accuracies could be higher with a smaller number of trials.

The motivation of this work is associated with BMIs, but these considerations apply to other neuroimaging studies that aim to characterize neural correlates with repetitive behavioral tasks. Recent approaches are also exploring the use of predictive models to relate neural features with cognitive states or neurological disorders^78–80^. In this context, models and cross-validation methods are used to confirm if a particular neural feature has some predictive power, which is then interpreted as an indication that the predictive feature is related to the cognitive variable of interest. Given that these studies are also likely to include long experimental sessions with a repetitive task, it is plausible that the neural signatures of interest could also change throughout the session and reduce the accuracies of the predictive models. Such reduced accuracies could yield interpretations that underestimate a neural feature’s role in the cognitive state of interest.

## Conclusion

In this study, we found that alpha- and beta-band ERDs associated with movement can change throughout trials in a repetitive motor task that is typically used in BMI studies. We also found that these changes can be associated with within session changes in classification performance in predicting when a participant is moving or resting. Further studies are needed to help identify how parameters of the behavioral tasks and an individual’s cognitive states can change movement-related ERDs over time and how these changes can be accounted for in the subsequent analyses. Our results also highlight the importance of reporting the number of trials and the time duration of the behavioral tasks when BMI’s predictive power is reported. These factors can significantly change neural modulations associated with repetitive tasks over time and are important to consider in neuroimaging studies that utilize predictive models to characterize brain activity and predict limb movements.

## Author Contributions

Andrew Paek: Conceptualization; data curation; formal analysis; funding acquisition; investigation; methodology; software; validation; visualization; writing – original draft; writing – review and editing. Shikha Prashad: Conceptualization; funding acquisition; methodology; resources; supervision; writing – review and editing.

## Acknowledgements

This work is supported by the National Science Foundation under Grant # EEC-2127509 administrated by the American Society for Engineering Education (ASEE).

## Data Availability Statement

Data are open source and available through the original authors as cited in the text.

### Appendix

**Table A1).** Classification Accuracies for each of the cross-validation conditions. The mean across participants is shown for each cell, while the standard deviation across participants is shown in parenthesis.

|  | Luciw et al. |  |  | Schwarz et al. |  |  | Jeong et al. |  |  |
| --- | --- | --- | --- | --- | --- | --- | --- | --- | --- |
|  | Combined | Early | Late | Combined | Early | Late | Combined | Early | Late |
| **Alpha-Bi** | 65.8%<br>(9.4%) | 68.7%<br>(9.0%) | 63.3%<br>(10.8%) | 69.3%<br>(13.1%) | 68.0%<br>(16.8%) | 68.9%<br>(15.7%) | 63.5%<br>(7.1%) | 62.1%<br>(9.8%) | 62.3%<br>(6.7%) |
| **Alpha-Contra** | 62.1%<br>(8.8%) | 62.3%<br>(7.4%) | 60.1%<br>(12.1%) | 62.7%<br>(9.6%) | 61.9%<br>(13.2%) | 68.2%<br>(11.4%) | 61.2%<br>(7.3%) | 60.4%<br>(9.1%) | 61.7%<br>(8.6%) |
| **Alpha-Ipsi** | 64.0%<br>(7.8%) | 67.5%<br>(9.0%) | 60.9%<br>(10.0%) | 69.8%<br>(12.7%) | 68.9%<br>(15.2%) | 66.5%<br>(14.5%) | 62.1%<br>(5.8%) | 61.0%<br>(9.5%) | 59.9%<br>(6.5%) |
| **Beta-Bi** | 73.4%<br>(7.5%) | 74.8%<br>(9.1%) | 77.2%<br>(8.3%) | 77.0%<br>(10.6%) | 78.1%<br>(12.3%) | 78.0%<br>(13.9%) | 66.5%<br>(8.7%) | 67.7%<br>(9.7%) | 64.8%<br>(8.8%) |
| **Beta-Contra** | 72.3%<br>(6.2%) | 69.4%<br>(7.6%) | 75.9%<br>(8.0%) | 73.0%<br>(9.4%) | 72.4%<br>(10.4%) | 74.0%<br>(12.1%) | 64.9%<br>(8.0%) | 65.6%<br>(9.0%) | 63.8%<br>(8.7%) |
| **Beta-Ipsi** | 68.9%<br>(8.8%) | 69.7%<br>(9.6%) | 69.6%<br>(9.2%) | 73.3%<br>(11.8%) | 73.9%<br>(12.0%) | 74.6%<br>(14.0%) | 61.5%<br>(8.2%) | 63.9%<br>(10.1%) | 60.2%<br>(8.2%) |

Early, trials from the first third of the experiment included; Late, trials from the last third of the experiment included; Combined, both early and late trials included; Bi, bilateral sensors; Contra, sensors contralateral to the engaged limb; Ipsi, sensors ipsilateral to the engaged limb.

**Table A2).**
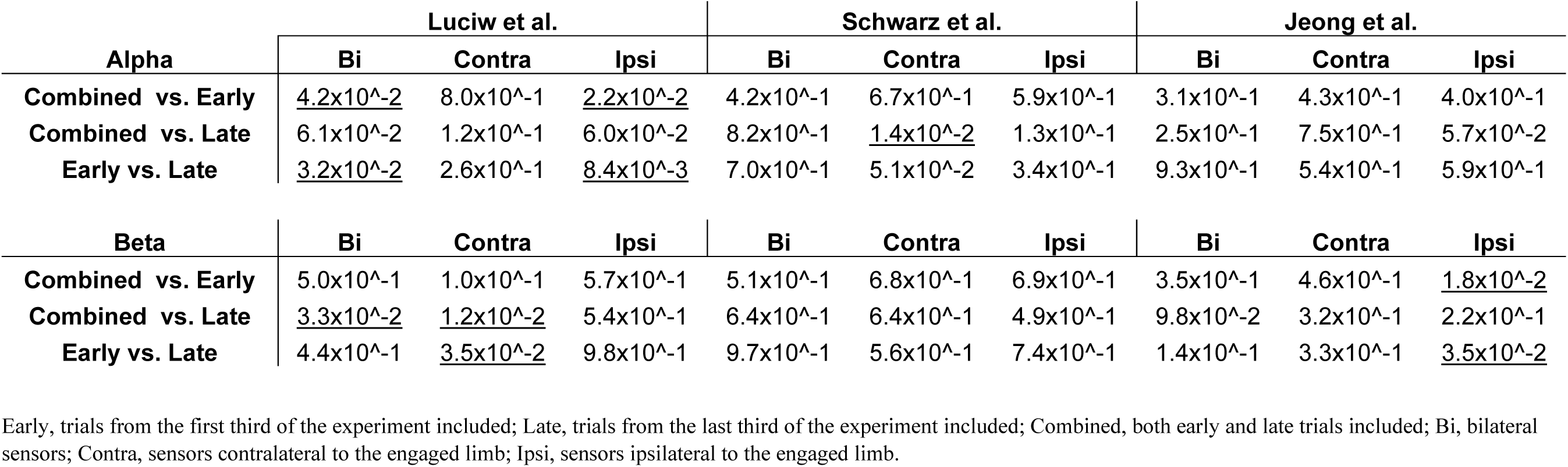
P-values of the t-test pair wise comparisons in classification accuracies between the trial-group permutations. Underlined p-values indicate significant differences (p < .05).

**Table A3).** P-values of the t-test pair wise comparisons in classification accuracies between groups of channels. Underlined p-values indicate significant differences (p < .05).

| Alpha | Luciw et al. |  |  | Schwarz et al. |  |  | Jeong et al. |  |  |
| --- | --- | --- | --- | --- | --- | --- | --- | --- | --- |
|  | Bi vs Con | Bi vs Ipsi | Con vs. Ipsi | Bi vs Con | Bi vs Ipsi | Con vs. Ipsi | Bi vs Con | Bi vs Ipsi | Con vs. Ipsi |
| Combined | <u>$6.7 \times 10^{-3}$</u> | $1.1 \times 10^{-1}$ | $1.9 \times 10^{-1}$ | <u>$1.6 \times 10^{-3}$</u> | $6.0 \times 10^{-1}$ | <u>$3.6 \times 10^{-3}$</u> | <u>$3.5 \times 10^{-3}$</u> | $1.5 \times 10^{-1}$ | $5.2 \times 10^{-1}$ |
| Early | <u>$2.8 \times 10^{-3}$</u> | $1.1 \times 10^{-1}$ | <u>$1.4 \times 10^{-2}$</u> | <u>$3.9 \times 10^{-2}$</u> | $3.9 \times 10^{-1}$ | <u>$9.5 \times 10^{-3}$</u> | $1.8 \times 10^{-1}$ | $2.0 \times 10^{-1}$ | $7.0 \times 10^{-1}$ |
| Late | <u>$1.0 \times 10^{-2}$</u> | $1.0 \times 10^{-1}$ | $6.2 \times 10^{-1}$ | $6.0 \times 10^{-1}$ | $1.6 \times 10^{-1}$ | $3.6 \times 10^{-1}$ | $5.2 \times 10^{-1}$ | <u>$1.2 \times 10^{-2}$</u> | $2.6 \times 10^{-1}$ |
| Beta | Bi vs Con | Bi vs Ipsi | Con vs. Ipsi | Bi vs Con | Bi vs Ipsi | Con vs. Ipsi | Bi vs Con | Bi vs Ipsi | Con vs. Ipsi |
|  | Bi vs Con | Bi vs Ipsi | Con vs. Ipsi | Bi vs Con | Bi vs Ipsi | Con vs. Ipsi | Bi vs Con | Bi vs Ipsi | Con vs. Ipsi |
| Combined | $2.2 \times 10^{-1}$ | <u>$1.1 \times 10^{-3}$</u> | <u>$1.6 \times 10^{-2}$</u> | <u>$7.3 \times 10^{-3}$</u> | <u>$1.3 \times 10^{-2}$</u> | $9.2 \times 10^{-1}$ | $7.4 \times 10^{-2}$ | <u>$7.9 \times 10^{-4}$</u> | $5.9 \times 10^{-2}$ |
| Early | <u>$1.3 \times 10^{-4}$</u> | <u>$8.4 \times 10^{-5}$</u> | $8.5 \times 10^{-1}$ | <u>$4.1 \times 10^{-2}$</u> | <u>$3.0 \times 10^{-2}$</u> | $6.3 \times 10^{-1}$ | $6.6 \times 10^{-2}$ | <u>$2.4 \times 10^{-2}$</u> | $4.6 \times 10^{-1}$ |
| Late | $1.8 \times 10^{-1}$ | <u>$3.2 \times 10^{-4}$</u> | <u>$2.9 \times 10^{-4}$</u> | $6.4 \times 10^{-2}$ | $1.6 \times 10^{-1}$ | $8.5 \times 10^{-1}$ | $3.9 \times 10^{-1}$ | <u>$4.3 \times 10^{-3}$</u> | $6.7 \times 10^{-2}$ |

Early, trials from the first third of the experiment included; Late, trials from the last third of the experiment included; Combined, both early and late trials included; Bi, bilateral sensors; Contra, sensors contralateral to the engaged limb; Ipsi, sensors ipsilateral to the engaged limb.

**Table A4).**
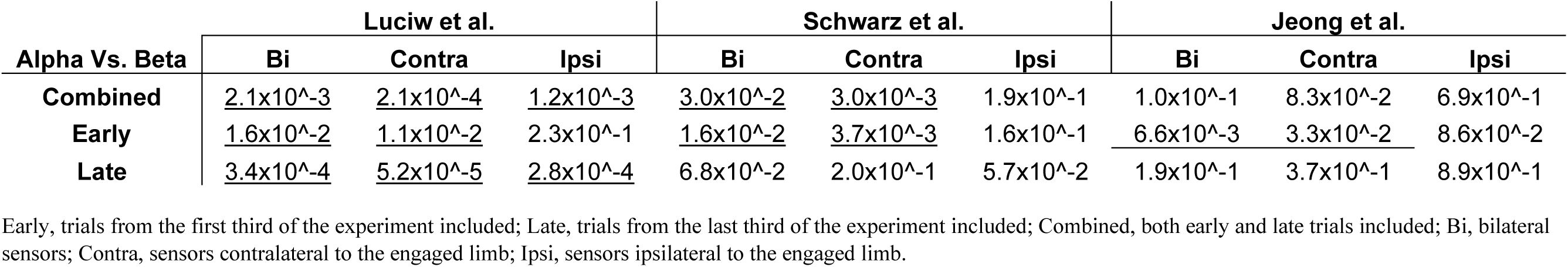
P-values of the t-test pairwise comparisons in classification accuracies between using alpha or beta band as the predictive feature. Underlined p-values indicate significant differences (p < .05).

| Alpha Vs. Beta | Luciw et al. |  |  | Schwarz et al. |  |  | Jeong et al. |  |  |
| --- | --- | --- | --- | --- | --- | --- | --- | --- | --- |
|  | Bi | Contra | Ipsi | Bi | Contra | Ipsi | Bi | Contra | Ipsi |
| **Combined** | <u>$2.1 \times 10^{-3}$</u> | <u>$2.1 \times 10^{-4}$</u> | <u>$1.2 \times 10^{-3}$</u> | <u>$3.0 \times 10^{-2}$</u> | <u>$3.0 \times 10^{-3}$</u> | $1.9 \times 10^{-1}$ | $1.0 \times 10^{-1}$ | $8.3 \times 10^{-2}$ | $6.9 \times 10^{-1}$ |
| **Early** | <u>$1.6 \times 10^{-2}$</u> | <u>$1.1 \times 10^{-2}$</u> | $2.3 \times 10^{-1}$ | <u>$1.6 \times 10^{-2}$</u> | <u>$3.7 \times 10^{-3}$</u> | $1.6 \times 10^{-1}$ | <u>$6.6 \times 10^{-3}$</u> | <u>$3.3 \times 10^{-2}$</u> | $8.6 \times 10^{-2}$ |
| **Late** | <u>$3.4 \times 10^{-4}$</u> | <u>$5.2 \times 10^{-5}$</u> | <u>$2.8 \times 10^{-4}$</u> | $6.8 \times 10^{-2}$ | $2.0 \times 10^{-1}$ | $5.7 \times 10^{-2}$ | $1.9 \times 10^{-1}$ | $3.7 \times 10^{-1}$ | $8.9 \times 10^{-1}$ |

Early, trials from the first third of the experiment included; Late, trials from the last third of the experiment included; Combined, both early and late trials included; Bi, bilateral sensors; Contra, sensors contralateral to the engaged limb; Ipsi, sensors ipsilateral to the engaged limb.

